# Indexing sensory plasticity: Evidence for distinct Predictive Coding and Hebbian Learning mechanisms in the cerebral cortex

**DOI:** 10.1101/189944

**Authors:** M. J. Spriggs, R. L. Sumner, R. L. McMillan, R. J. Moran, I. J. Kirk, S. D. Muthukumaraswamy

**Affiliations:** School of Psychology, The University of Auckland; School of Pharmacy, The University of Auckland; Brain Research New Zealand; Department Engineering Mathematics, University of Bristol BS8 1TH, UK

**Author notes:** M.J.S and R.L.S contributed equally to this work. Corresponding Author: Meg J Spriggs, School of Psychology, University of Auckland., Private Bag 92019, Auckland 1142, New Zealand.

**Keywords:** Dynamic Causal Modelling, Long Term Potentiation, Mismatch Negativity, Neuroplasticity, Perceptual Learning

## Abstract

- ERP and DCM study of two sensory plasticity paradigms: roving MMN and visual LTP
- First demonstration of multiple learning mechanisms under different task demands
- Evidence for both Predictive Coding and Hebbian learning mechanisms
- The BDNF Val66Met polymorphism modulates ERPs for both paradigms
- However, the polymorphism only modulates MMN network connectivity

The Roving Mismatch Negativity (MMN), and Visual LTP paradigms are widely used as independent measures of sensory plasticity. However, the paradigms are built upon fundamentally different (and seemingly opposing) models of perceptual learning; namely, Predictive Coding (MMN) and Hebbian plasticity (LTP). The aims of the current study were to 1) compare the generative mechanisms of the MMN and visual LTP, therefore assessing whether Predictive Coding and Hebbian mechanisms co-occur in the brain, and 2) assess whether the paradigms identify similar group differences in plasticity. Forty participants were split into two groups based on the *BDNF* Val66Met polymorphism and were presented with both paradigms. Consistent with Predictive Coding and Hebbian predictions, Dynamic Causal Modelling revealed that the generation of the MMN modulates forward and backward connections in the underlying network, while visual LTP only modulates forward connections. Genetic differences were identified in the ERPs for both paradigms, but were only apparent in backward connections of the MMN network. These results suggest that both Predictive Coding and Hebbian mechanisms are utilized by the brain under different task demands. Additionally, both tasks provide unique insight into plasticity mechanisms, which has important implications for future studies of aberrant plasticity in clinical populations.

## 1. Introduction

Perceptual learning relies on the structural and functional modification of neural networks in response to external stimulation (Fahle, 2004). This experience-dependent neuroplasticity within the sensory systems provides an opportunity to non-invasively study the mechanisms underlying neuroplasticity throughout the brain. However, different external demands (e.g., task demands) may elicit different encoding mechanisms (Koch & Poggio, 1999) and to date, the differences between such mechanisms have not been characterized.

The most widely studied form of neuroplasticity is Long Term Potentiation (LTP). LTP refers to an activity dependent increase in synaptic efficacy following repeated neuronal co-activation, and is dependent on an influx of Ca^2+^ through N-methyl-D-aspartate receptors (NMDAR) leading to long term alterations in cell structure and function (Bliss & Lømo, 1973; Cooke & Bliss, 2006; Teyler & DiScenna, 1987). Importantly, LTP conforms to many Hebbian characteristics such as input-specificity, co-activation and associativity (Hebb, 1949). As such, Hebbian LTP is regarded as the most likely neuronal mechanism underlying memory formation.

LTP has been primarily studied in laboratory animals using direct neuronal electrical stimulation (Bliss & Lømo, 1973; Figurov, Pozzo-Miller, Olafsson, & others, 1996; Harris, Ganong, & Cotman, 1984; Kirkwood & Bear, 1994; Teyler & DiScenna, 1987). However, following the demonstration of visually-induced enhancements in the neural activation of rodents (Heynen & Bear, 2001), Teyler et al., (2005) presented one of the first electroencephalography (EEG) paradigms for measuring LTP-*like* mechanisms noninvasively in humans. High frequency (~9Hz) visual stimulation was used to induce an enhancement of the visually evoked potential (VEP) to later low frequency (~1Hz) presentations of the same stimulus. Subsequent human and rodent studies have demonstrated that this visually-induced enhancement conforms to many of the Hebbian characteristics seen in rodent LTP such as longevity, NMDAR dependence (Clapp, Eckert, Teyler, & Abraham, 2006) and input specificity (McNair et al., 2006; Ross et al., 2008). Furthermore, this paradigm has been used to demonstrate modulated plasticity in healthy and clinical populations (Çavuş et al., 2012; Normann, Schmitz, Fürmaier, Döing, & Bach, 2007; Smallwood et al., 2015; Spriggs, Cadwallader, Hamm, Tippett, & Kirk, 2017). Together, this body of human and rodent studies indicates that this visually induced enhancement represents the induction of a Hebbian LTP-*like* form of neuroplasticity (Clapp, Hamm, Kirk, & Teyler, 2012; Kirk et al., 2010).

While potentiation of the VEP has been well characterized, modulations to the underlying network remain largely unexplored. Both EEG source localization and functional magnetic resonance imaging (fMRI) have localized the LTP-*like* enhancement to extrastriate visual cortex (Clapp et al., 2005; Teyler et al., 2005). From extrastriate visual cortex, the ventral and dorsal visual streams extend to the medial temporal lobe and parietal lobe respectively (Felleman & Van Essen, 1991; Grill-Spector & Malach, 2004). Experience-dependent plasticity within these networks is understood to underlie visual perceptual learning (Fahle, 2004; Kourtzi & DiCarlo, 2006), with changes occurring at some of the earliest levels of cortical processing (Cooke & Bear, 2014; Kourtzi & DiCarlo, 2006). The ventral visual stream is understood to support object recognition, and is closely intertwined with medial temporal memory networks (Desimone et al., 1985; Felleman & Van Essen, 1991; Grill-Spector & Malach, 2004; Kourtzi & DiCarlo, 2006). As such, one can speculate that LTP-induction will enhance connectivity within this ventral visual network.

The characteristics of Hebbian plasticity provide a framework for long-term, experience-dependent enhancements in synaptic responses. However, Hebbian plasticity is not the only mechanism for experience-dependent plasticity in the neocortex, and growing emphasis is being placed on Bayesian models of perceptual learning. Such models propose that the brain is equipped with a generative model, which is built upon prior expectations extracted from sensory data and provides a mapping of (hidden) cause to (sensory) consequence (Friston, 2005; Knill & Pouget, 2004). The Predictive Coding model proposes that prediction errors are used to adjust the generative model until divergence is minimized; allowing for an accurate model of the cause of incoming information (Bastos et al., 2012; Friston, 2005; Garrido, Kilner, Stephan, & Friston, 2009a; Huang & Rao, 2011). The reduction of prediction error is dependent on the passing of top down predictions and bottom up prediction errors through hierarchical, reciprocally connected networks. Neurocomputational modelling of prediction errors suggests that top-down predictions are expressed through NMDAR and GABAergic pathways, while bottom up prediction errors rely on fast feedback via AMPA receptors (Corlett, Honey, & Fletcher, 2016). Under the Predictive Coding framework, experience-dependent plasticity corresponds to the reciprocal updating of internal models of the environment through these pathways.

The most studied empirical example of Predictive Coding in the brain is the Mismatch Negativity (MMN). The MMN is a large, fronto-central negativity induced by a surprising or ‘deviant’ tone following a sequence of predictable or ‘standard’ tones, (Garrido, Kilner, Stephan, et al., 2009). The widely used ‘roving MMN’ paradigm involves the presentation of trains of tones of the same frequency, where the first (deviant) tone in each train induces the MMN response, and this subsequently returns to a standard response over successive presentations. Under the predictive coding framework, the MMN represents a failure to predict bottom-up sensory input and, consequently to suppress prediction error (Friston, 2005; Garrido, Kilner, Stephan, et al., 2009). In support of this, previous studies have demonstrated that the MMN is generated by modulations in intrinsic auditory cortex (A1) connectivity, as well as reciprocal message passing within a fronto-temporal network (Auksztulewicz & Friston, 2015; Garrido et al., 2008; Garrido, Kilner, Kiebel, Stephan, & Friston, 2007; Moran, Symmonds, Dolan, & Friston, 2014; Schmidt et al., 2013). This suppression of prediction error corresponds to perceptual inference (Auksztulewicz & Friston, 2016; Garrido, Kilner, Stephan, & Friston, 2009). The MMN paradigm has been used to demonstrate disrupted perceptual inference in clinical populations (Boly et al., 2011; Dima, Dietrich, Dillo, & Emrich, 2010) and under pharmacological intervention (Rosch, Auksztulewicz, Leung, Friston, & Baldeweg, 2017; Schmidt et al., 2013).

While investigating the MMN alone indexes plasticity over peri-stimulus time, the roving MMN paradigm allows additional exploration of changes in the mismatch response over successive trials. Under Predictive Coding, repetition-dependent changes are a means by which the generative model is optimized to provide a more accurate and precise model of sensory inputs. The optimization of the model parameters (and the precision of predictions) across repetitions corresponds to perceptual learning (Auksztulewicz & Friston, 2016; Garrido, Kilner, Stephan, et al., 2009). These precision parameters are mediated by a number of different neuromodulators including dopamine, acetylcholine, and serotonin (Corlett et al., 2016; Friston, Brown, Siemerkus, & Stephan, 2016; Moran et al., 2013). Pharmaco-EEG studies using Ketamine (a NMDA antagonist at low doses) have also demonstrated the strong relationship between the balance of NMDAR signaling and the repetition effects (Rosch et al., 2017; Schmidt et al., 2013). Ketamine, has been found to modulate NMDA‐ and AMPA-mediated frontal-to-parietal connectivity as well as NMDA-mediated GABAergic inhibitory interneuronal connectivity within frontal microcircuits (Muthukumaraswamy et al., 2015; Rosch et al., 2017). Studies employing the roving MMN paradigm have also have demonstrated that repetition dependent changes in intrinsic (precision) and extrinsic (prediction error) connectivity occur very quickly, implicating short-term plasticity mechanisms (Garrido, Kilner, Stephan, et al., 2009; Rosch et al., 2017; Schmidt et al., 2013).Together, the roving MMN paradigm can thus be used as an index of both perceptual inference (the MMN itself) and perceptual learning (repetition suppression) which jointly index sensory plasticity under a Predictive Coding framework.

As illustrated above, both Hebbian and Predictive Coding mechanisms have been independently implicated in perceptual learning and the visual LTP and roving MMN paradigms were designed to index these models respectively. However, the two models are built upon fundamentally different assumptions of how perceptual learning is encoded in the brain; primarily, while Predictive Coding is dependent on updating an internal, generative model, Hebbian plasticity is not. The coexistence of Hebbian and Predictive Coding mechanisms has been explored in models of cortical responses such as the Free Energy Principle (Friston, 2005, 2009, 2010). Under the Free Energy Principle, Predictive Coding and Hebbian mechanisms are used to define hidden states and causes of an internal generative model respectively (Bastos et al., 2012; Friston, 2010). However, it may be possible that Hebbian processes can occur independent of a generative model, and that the brain may employ different encoding mechanisms for different tasks (Koch & Poggio, 1999). As such, the first aim of the current study was to compare the mechanisms underlying the generation of the MMN using the roving MMN paradigm, and the potentiated VEP using the visual LTP paradigm. It was hypothesized that the paradigms would induce different changes within the underlying neural network. Specifically, as the primary difference between Hebbian and Predictive Coding models is dependence on a generative model, it was hypothesized that the paradigms would differ in their modulation of top-down connectivity.

An important assumption of studies using these paradigms with clinical populations is that they are sensitive to aberrant plasticity. However, under the hypothesis that the brain uses both Predictive Coding and Hebbian mechanisms in different circumstances, it is unclear whether aberrant plasticity will manifest uniformly across these encoding mechanisms. As such, a secondary aim of the current study was to assess the consistency of group differences across the two paradigms. Participants were split into two groups based on the *BDNF* Val66Met polymorphism. The *BDNF* gene codes for Brain Derived Neurotrophic Factor; an important molecular mediator of neural plasticity (Goldberg & Weinberger, 2004; Park & Poo, 2013; Tyler, Alonso, Bramham, & Pozzo-Miller, 2002). Approximately 25-50% of the population carry the rs6265 single nucleotide polymorphism (SNP) (4-16% homozygous) which substitutes valine to methionine at codon 66 (known as Val66Met) (Goldberg & Weinberger, 2004; Shimizu, Hashimoto, & Iyo, 2004). The Met allele of the SNP is associated with reduced secretion of BDNF, and thus has previously been implicated in the efficacy of NMDAR-dependent neuroplasticity and memory performance (Chen et al., 2004; Egan et al., 2003; Hariri et al., 2003; Lamb et al., 2015). Previous studies conducted in our lab using the LTP paradigm have demonstrated reduced potentiation of the N1b component of the VEP in *BDNF* Met carriers (Thompson et al., *in prep*). Soltész et al., (2014) assessed the effect of the Val66Met polymorphism on the MMN and found no effect on MMN amplitude. There was however, no analysis of repetition suppression. Because repetition suppression is NMDAR mediated, it is reasonable to hypothesize that there may be an effect of the Val66Met polymorphism. Furthermore, neither of the above studies assessed genotype differences in connectivity modulation in the networks generating the ERPs, and it was hypothesized that group differences would also be apparent in the modulation of connectivity by the two paradigms.

## 2. Materials and Methods

### 2.1. Participants

44 male and female participants volunteered for the study (age range: 19-33, 33 female and 7 male; the imbalance in gender split is due to overlap of participants with another study). Four participants were excluded from the final analysis due to insufficient data quality, leaving a final sample of 40. Participants were required to have no history of neurological conditions or concussion, and normal or corrected to normal vision. This study was approved by the University of Auckland Human Participants Ethics Committee. Participants provided informed written consent prior to participation.

### 2.2. Genotyping

Saliva samples were collected using Oragene Self Collection kits. DNA was extracted and genotyped for *BDNF* Val66Met by the Auckland Sequenom Facility using the TaqMan 5’-exonuclease assay. Primers and TaqMan probes were from Applied Biosystems Inc. Electrophoresis was used to determine the genotype for each participant. From this, participants were split into two groups defined by *BDNF* Val66Met genotype: Val homozygotes (Val/Val *N* = 21), and Met carriers (Val/Met *N*= 13, Met/Met *N*= 6). The grouping of Met homozygotes and heterozygotes into a single group is due to the low prevalence of Met homozygotes in the general population (4-16%, Shimizu et al., 2004), and is consistent with a large body of previous literature (Kambeitz et al., 2012).

### 2.3. Equipment

EEG data were collected using 64 channel Acticap Ag/AgCl active shielded electrodes and Brain Products MRPlus amplifiers recorded in Brain Vision Recorder (Brain Products GmbH, Germany) with a 1000Hz sampling rate, and 0.μV resolution. FCz was used as an online reference, AFz as ground. Electrode impedance was maintained below 25kΩ.

Stimuli were displayed on an ASUS VG248QE computer monitor with a screen resolution of 1920 x 1800 and 144Hz refresh rate. TTL pulses generated through the parallel port of the display computer provided synchronisation of stimulus events with EEG acquisition.

### 2.4. Tasks

All participants were presented with both the MMN and LTP task. To avoid carry-over effects, the presentation order was such that for 25% of participants the MMN task preceded the LTP task, for 25% it followed the LTP task, and for 50% it took place during the rest period of the LTP task.

#### 2.4.1. Mismatch negativity

EEG was recorded continuously while participants engaged in a roving auditory oddball task used to probe the mismatch negativity in response to unattended stimuli (Figure 1i; Garrido et al., 2008). The task was written and run in MATLAB using the Cogent toolbox (www.vislab.ucl.ac.uk/cogent.php).

**Figure 1.**
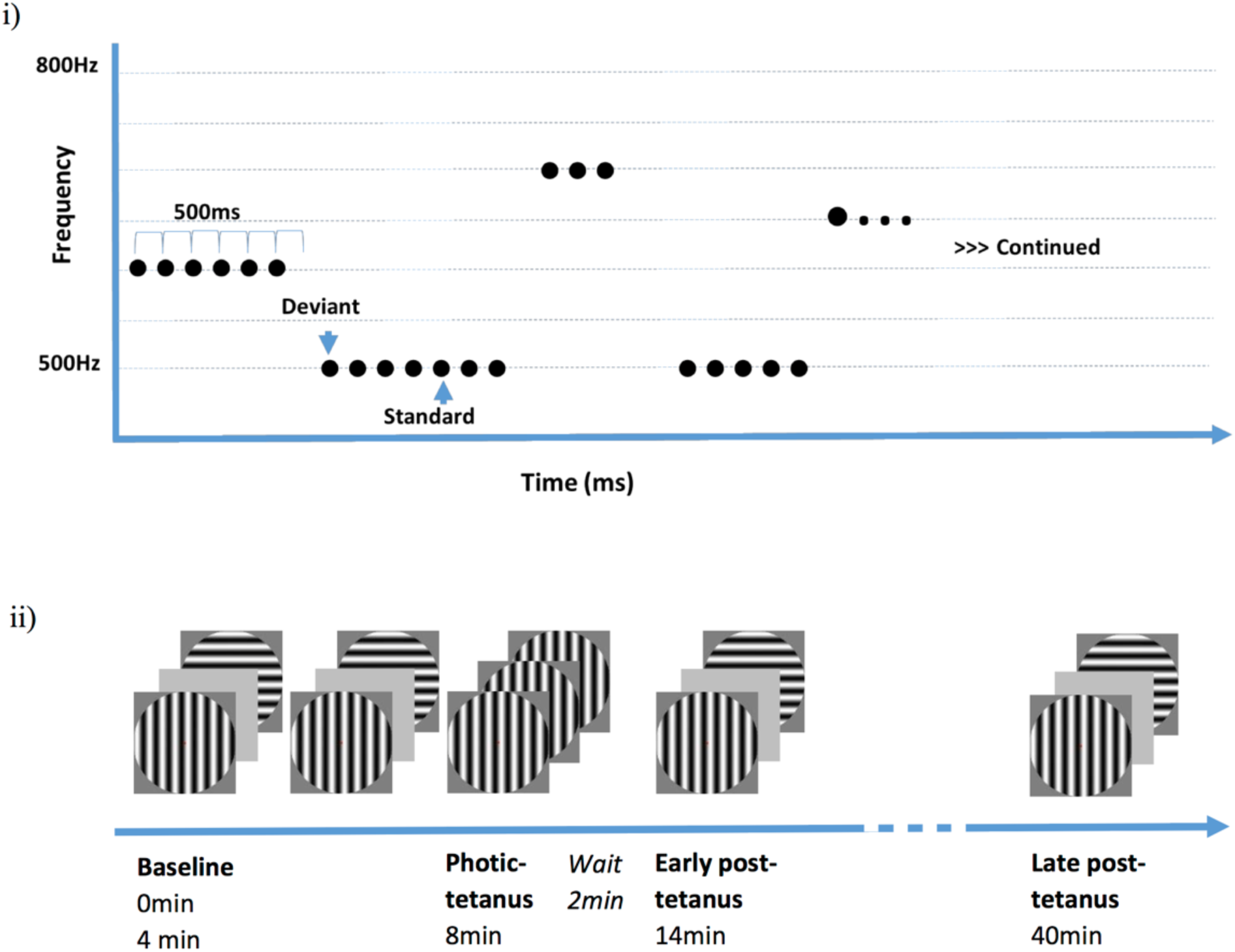
i) Depiction of the Roving MMN paradigm where the first and sixth tone in each sequence act as the deviant and standard respectively, ii) The condition structure and timing for the LTP task.

The stimuli consisted of trains of one to eleven identical sinusoidal tones. The first tone of each train was treated as the deviant tone (typically producing the classic “oddball” response), while tone 6 was treated as standard. As such, the oddball and standard in a given train have the exact same physical properties, differing only in the number of preceding presentations. The variability in the number of tones in a train prevented higher order regularity (such as change anticipation). Pseudo-randomisation of train length produced 250 deviant presentations of which, 2.5% had one or no repetitions; 3.75% had 2 and 3 repetitions; and 12.5% had 5 to 10 repetitions.

Tone frequency varied within 500 and 800Hz in random steps of integer multiples of 50Hz. The tones were 70ms in length with a 5ms rise and fall time, and 500ms inter-stimulus interval (ISI). Tones were presented binaurally at a constant volume that was adjusted for individual participants so that it was clear, and comfortable.

Participants were instructed to focus on a visual distractor task, where they were required to press the spacebar key when they detected a change in stimulus luminance. The stimulus was a small fixation cross that changed luminance pseudo-randomly every 2-5 seconds (unrelated to auditory changes). The change in luminance appeared as a somewhat subtle change between black and grey and therefore demanded substantial attention from the participant.

#### 2.4.2. Visual LTP

Sensory LTP was measured using a slight modification of an established paradigm for inducing LTP-*like* enhancements of early VEP components (Figure 1ii; McNair et al., 2006; Teyler et al., 2005). The task was written and run in MATLAB, using the Psychophysics Toolbox (Brainard, 1997; Kleiner et al., 2007; Pelli, 1997), with a gamma correction applied to the screen.

The stimuli used were circular vertical and horizontal sine gratings with a spatial frequency of 1 cycle per degree. Stimuli were presented at full contrast on a grey background subtending 8 degrees of visual angle. For all conditions participants were seated with their eyes 90cm from the centre of the screen and were instructed to passively fixate on a centrally presented red dot.

The task comprised four conditions. For the first condition (referred to hereafter as pre-tetanus), both stimuli were presented in a random order 240 times (480 presentations in total) for 34.8ms at a temporal frequency of 1Hz. The interstimulus interval was varied using 5 intervals from 897-1036ms that occurred randomly but equally often. This condition took approximately 8 minutes.

The second condition was the photic tetanus or high frequency stimulation, and directly followed the pre-tetanus condition. This consisted of 1000 presentations of either the horizontal or vertical stimulus (counterbalanced between participants) for 34.8ms with a temporal frequency of approximately 9Hz. The interstimulus interval was either 62.6 or 90.4ms occurring at random but equally often. This condition took approximately 2 minutes.

The third condition was an early post-tetanus condition that followed 2 minutes after the tetanus, allowing retinal after images to dissipate. The fourth condition was a late post-tetanus block that took place 30mins after the early post-tetanus condition. Both the post tetanus conditions had the same parameters as the pre-tetanus condition, but were split across the two time points (240 trials each block as opposed to 480) therefore establishing the change in response from baseline both immediately following high frequency stimulation, and after an extended break. Each post-tetanus condition took approximately 4 minutes.

### 2.5. Data collection and preprocessing

All preprocessing and data analyses were performed using SPM12 (http://www.fil.ion.ucl.ac.uk/spm/software/spm12/). Data were downsampled to 250Hz and re-referenced to the common average. A 0.1-30Hz bandpass filter was used to remove both high and low frequency noise. Eye blinks were identified and removed by thresholding electro-oculogram (EOG) channels (or Fp1 and Fp2 when EOG channels were not available). Visual artifact rejection was performed using the FieldTrip visual artifact rejection tools in SPM.

#### 2.5.1. MMN

The MMN data were baseline corrected and segmented into 500ms epochs (-100 - 400ms). Trials were averaged based on tone repetition, collapsed across frequencies. This resulted in averaged responses for tone presentations 1-10. Tone 1 was treated as the deviant tone, and tone 6 was treated as the standard tone.

#### 2.5.2. LTP

The LTP data were baseline corrected and segmented in 600ms epochs (-100ms – 500ms). Data were then averaged based on stimulus condition (tetanized stimulus, non-tetanized stimulus) for each of the three time points independently (pre-tetanus, early post-tetanus, late post-tetanus).

### 2.6. Analysis of ERPs

For all analyses, main effects were considered significant at *p* < 0.05 family-wise error corrected (FWE-c). A more liberal threshold is reported for interaction effects (*p* < 0.001 uncorrected). Simple effects tests were conducted as appropriate. Where multiple significant peaks occur for the same component, a significant time window is reported, taken from the *t*-distribution and only the most significant peak within a cluster is reported.

#### 2.6.1. LTP

The preprocessed data were converted into NIfTI images using a time window of 0-250ms, and were smoothed using a 6 x 6 x 6 FWHM Gaussian kernel. An initial manipulation confirmation of the induction of potentiation was conducted by running a 3 x 2 ANOVA that probed the effects of time (pre-tetanus, early post-tetanus, and late post-tetanus) and stimulus type (tetanised and non-tetanised). In a subsequent analysis, a 2 x 2 x 2 ANOVA that probed the effect of time (early vs late tetanus), tetanus (tetanized vs non-tetanized), and genotype (Val homozygotes vs Met carriers) was run using difference waves of early post-tetanus block minus pre-tetanus and, late post-tetanus block minus pre-tetanus. An occipito-parietal ROI, informed by the prior analysis and by previous studies (McNair et al., 2006; Ross et al., 2008) was used, comprising the following electrodes: P1, P2, P3, P4, P5, P6, P7, P8, Pz, PO3, PO4, PO7, PO8, PO9, PO10, POz, O1, O2, and Oz.

#### 2.6.2. MMN

The preprocessed data were converted to NIfTI images using a time window of 0-400ms, and images were smoothed using a 6 x 6 x 6 FWHM Gaussian kernel. An initial *t*-test between deviant and standard (i.e., tones 1 and 6) was conducted to confirm elicitation of the MMN ERP. A two-tailed *t*-test was then run to compare the average amplitude of the MMN in response to the deviant tone in *BDNF* Val homozygotes and Met carriers. Finally, to assess tone repetition effects, difference waves were calculated between the deviant tone and subsequent tones up to the standard (i.e., tone 2, 3, 4, 5 and 6). A 2 x 5 ANOVA was then run that probed the effects of genotype (Val homozygotes, vs Met carriers) on these difference waves. Simple effects tests were conducted as appropriate.

### 2.7. Dynamic Causal Modelling

Dynamic Causal Modelling (DCM) was used to assess the network architecture underlying the generation of both the MMN and LTP ERPs. DCM uses biologically informed models within a Bayesian framework to infer hidden variables relating to the modulation of intrinsic and extrinsic connectivity by exogenous input (Friston, 2003). EEG data are modelled as perturbations in a non-linear, dynamic, input-state-output system where activity in one source is caused by activity in another. The generative model is comprised of a neuronal mass model and an electrophysiological model to determine how the hidden neuronal states translate to what is recorded on the scalp (David et al., 2006; Kiebel, David, & Friston, 2006). These biological constraints allow for neurobiologically plausible interpretations of ERPs as reflecting modulations of effective connectivity within a network.

#### 2.7.1. Model Specification

The neuronal mass modelling employed in the current study represents neuronal processes as the post-membrane potential and firing rate of three subpopulations of excitatory pyramidal cells, spiny stellate cells and inhibitory interneurons. The subpopulations are connected within each source via intrinsic connections, and between sources via extrinsic connections. Based on the Jansen and Rit model (Jansen & Rit, 1995) and the connectivity rules described by Felleman and Van Essen (1991), the directionality of extrinsic connections (forward, backward, or lateral) can be determined via the neuronal population from which they originate and terminate. These constraints allow for the construction of hierarchical cortico-cortical networks, and can be specified as a set of differential equations that describe the neural dynamics.

The neuronal model is then passed through the electrophysiological model, where each source is modelled as a single Equivalent Current Dipole (ECD). Lead field mapping is parameterized in terms of the location and orientation of each dipole (Kiebel et al., 2006). A four concentric sphere head model with homogeneous and isotropic conductivity is used as an approximation to the brain, cerebrospinal fluid, skull and scalp surfaces. The orientation parameters had a prior mean of zero, and a variance of 256mm^2^. For computational expediency, the sensor data in the current study were reduced to 8 dimensions by projecting the data onto a subspace defined by the principle eigenvectors (David et al., 2006).

The generative model is then inverted using a Variational Bayes scheme to assess parameter likelihood given the data and the model for each subject individually (Friston, 2002). This involves updating the posterior moments (mean and covariance) to minimize the free energy, *F*; an approximation to the log model evidence. This iterative procedure provides an approximation to the posterior probability of the model parameters *p*(θ\*y*,*m*), as well as an approximation to the model evidence *p*(*y*\*m*)used for model comparison.

#### 2.7.2. DCM statistics

Random effects (RFX) Bayesian Model Selection (BMS) was used to identify the model of best fit, using *F* as an approximation to model evidence. In the current study, the protected exceedance probability (PXP) was used as an index of model fit. The PXP quantifies the probability that any one model is more frequent than the others above and beyond chance (Rigoux, Stephan, Friston, & Daunizeau, 2014). The Bayesian Omnibus Risk (BOR) was then used as an index of the probability of having erroneously chosen *H*_1_ over *H*_0_. This is therefore the risk that the observed sample occurred by chance, which is comparable (though not equivalent) to a *p* value in classical statistics. BOR≈0.25 is considered strong evidence that there is a true difference in model frequencies (Rigoux et al., 2014).

Finally, *a posteriori* estimates of model parameters for the winning MMN and LTP models were then used for classical inference on parameter modulation by the paradigms. Specifically, estimates from individual parameters were subject to *t*-tests (p<.05, uncorrected).

#### 2.7.3. Source identification

Sources for the MMN and LTP paradigms were identified using group source inversion within the Multiple Sparse Priors method implemented in SPM12 (Litvak & Friston, 2008). The time windows for source localization were chosen for both paradigms based on the sensor space data. For the MMN, source inversion was performed on a 200-300ms time window post stimulus, which was then subject to a *t*-test comparing deviant (tone 1) to standard (tone 6) (p<.001 uncorrected). For the LTP paradigm, sources were identified in two time windows corresponding to the two time windows of interest: 128-132ms, and 188-208ms. For both time windows, source images were subject to 2(tetanus) x 3(time) ANOVAs and sources of interest were identified for the main effect of time (p<.05 uncorrected).

## 3. Results

### 3.1. MMN

#### 3.1.1. ERP analyses

*t*-tests comparing the standard tone ERP to the deviant tone revealed a significant difference between 210-310ms that peaked at 256ms (t_(39)_ = 10.45, *p* < 0. 001 FWE-c) (Figure 2). The *t*-distribution revealed a frontal cluster consistent with the MMN response. In addition, there was a left lateralized significant time-window of 348-380ms that peaked at 352ms (t_(39)_ = 4.57, *p* < 0.01 FWE-c).

**Figure 2.**
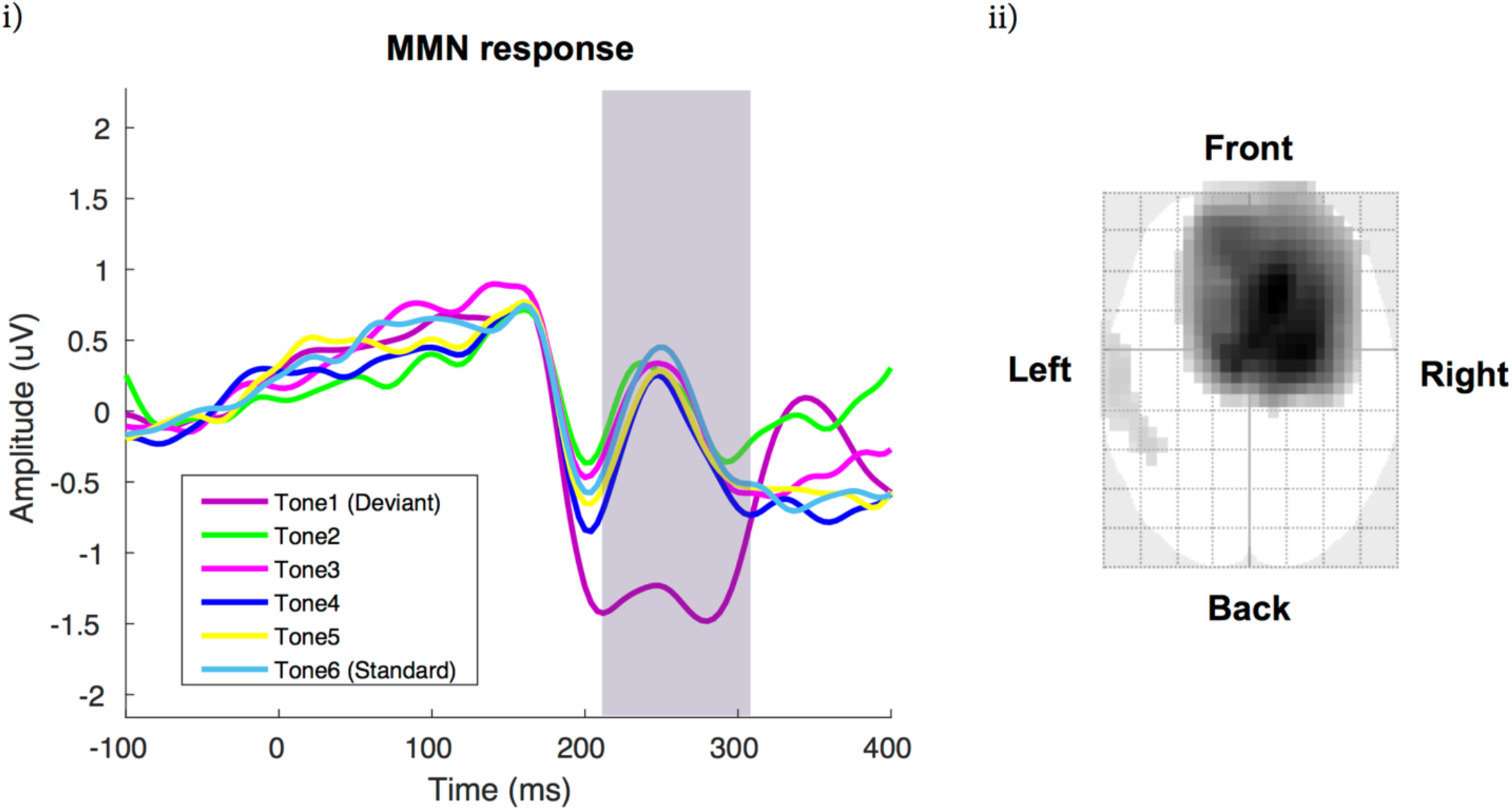
Averaged ERP across genotype i) the MMN response is significant from 210-310ms and is shown by the purple line (deviant) compared to the lightest blue line (standard). ii) The *t*-distribution of the significant MMN response (*p* < .05 FWE-c).

The *t*-test of genetic effects on the amplitude of the MMN response to the deviant tone revealed no significant differences (FWE-c).

To assess repetition suppression, a 2 x 5 ANOVA comparing the effect of genotype on the difference wave of the deviant tone minus subsequent tones (tone 2, 3, 4, 5, and 6) was performed. There was a significant main effect of tone number (see Supplementary Material) and a significant interaction between tone number and genotype.

Investigating an interaction between subsequent tone number and genotype probes the effect of genotype on variability between the difference waves for subsequent tones, thus exploring the progression in the ERP to a standard response. The interaction revealed a small but significant frontal cluster at 188ms that is consistent with the MMN (F_(4, 190)_ = 4.8, *p* = 0.001 uncorrected). A corresponding peak at 192ms (F_(4, 190)_ = 4.77, *p* = 0.001 uncorrected) was identified, which is consistent with the temporal positive deflection that has been reported to occur alongside the MMN (Rinne, Alho, Ilmoniemi, Virtanen, & Näätänen, 2000). An additional significant cluster (392-400ms) was identified that peaked at 392ms (F_(4, 190)_ = 4.72, *p* = 0.001 uncorrected). This was not found to be consistent with known ERPs in this time window such as the P3a and may reflect a late occipital positivity reported in some MMN studies (Auksztulewicz & Friston, 2015).

*Post-hoc* contrasts were performed that probed the interaction between genotype and the variability between each difference wave and it’s directly subsequent neighbor (e.g. tone 2 difference wave compared to tone 3 difference wave, tone 3 difference wave to tone 4 difference wave and so on). The purpose of this approach (as opposed to simply comparing the amplitude of the response to a tone) is to capture the progression from high variability (indicating a prediction error or deviant response) to no significant variability (indicating a standard response has developed) between successive tones. In doing so, this analysis examines the influence of genetic group on repetition suppression by comparing the time-course of the return to standard responding, with a sharper reduction in variability indicating more rapid learning of the new tone. An approach that analysed between tone variability, rather than simply measuring differences in amplitude between genotypes at each tone, was used because of the non-linear fashion with which the ERP returns to a standard amplitude (Figure 2).

From these *post-hoc* contrasts, a comparison of tone 2 to tone 3 between genotypes revealed a significant interaction; an occipital cluster from 368-400ms that peaked at 388ms (F_(4, 190)_ = 15.16, *p* < 0.001 uncorrected). This was attributed to the occipital component reported above. A contrast of tone 3 compared to tone 4 between genotypes revealed a frontal significant interaction from 188-215ms, peaking at 188ms (F_(4, 190)_ = 17.54, *p* < 0.001 uncorrected) consistent with the MMN (Figure 3 and Supplementary Material Figure S1). Within the same time-window the corresponding temporal positivity also appeared and peaked at 192ms (F_(4, 190)_ = 16.00, *p* < 0.001 uncorrected). This interaction is interpreted as greater variability for Val homozygotes (Figure 3i) compared to Met carriers (Figure 3ii) in the repetition effect of the MMN. Also within this interaction the occipital time-window of 384-400ms appeared and peaked at 384ms (F_(4, 190)_ = 14.68, *p* < 0.001 uncorrected). There were no significant peaks from tones 4-5. There was one frontal left lateralized small peak from tone 5-6 at 380ms (F_(4, 190)_ = 11.75, *p* < 0.001 uncorrected).

**Figure 3.**
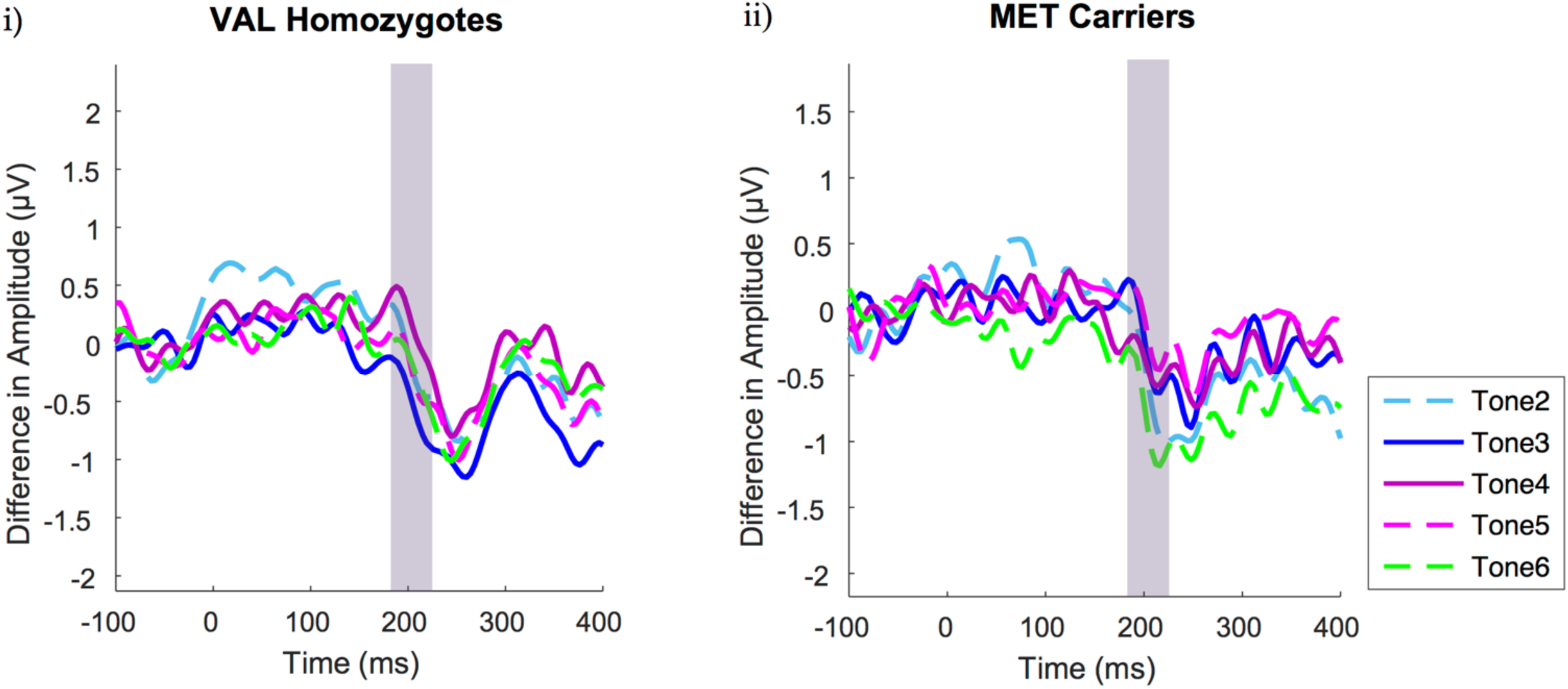
The difference between the deviant tone and each subsequent tone at electrode F5 illustrates the greater variability for Val homozygotes (i) compared to Met carriers (ii). The solid lines draw attention to the interaction capturing the difference between tone 3 and 4 compared to standard for each genotype (188-215ms). This indicates Met carriers produce a standard response sooner than Val homozygotes.

#### 3.1.2. Sources

Source analysis in the current study (Figure 4ii) revealed bilateral sources in STG (MNI coordinates left: -62,-30, 16, right: 52, -32, 6) and right IFG (MNI coordinates 48, 30 12) (*p*<.001, uncorrected). Coordinates for A1 sources were taken from Garrido et al., (2008) (MNI coordinates left: -42, -22, 7, right: 46, -14, 8). The locations for all 5 sources were used for subsequent dynamic causal modelling of the evoked responses.

**Figure 4.**
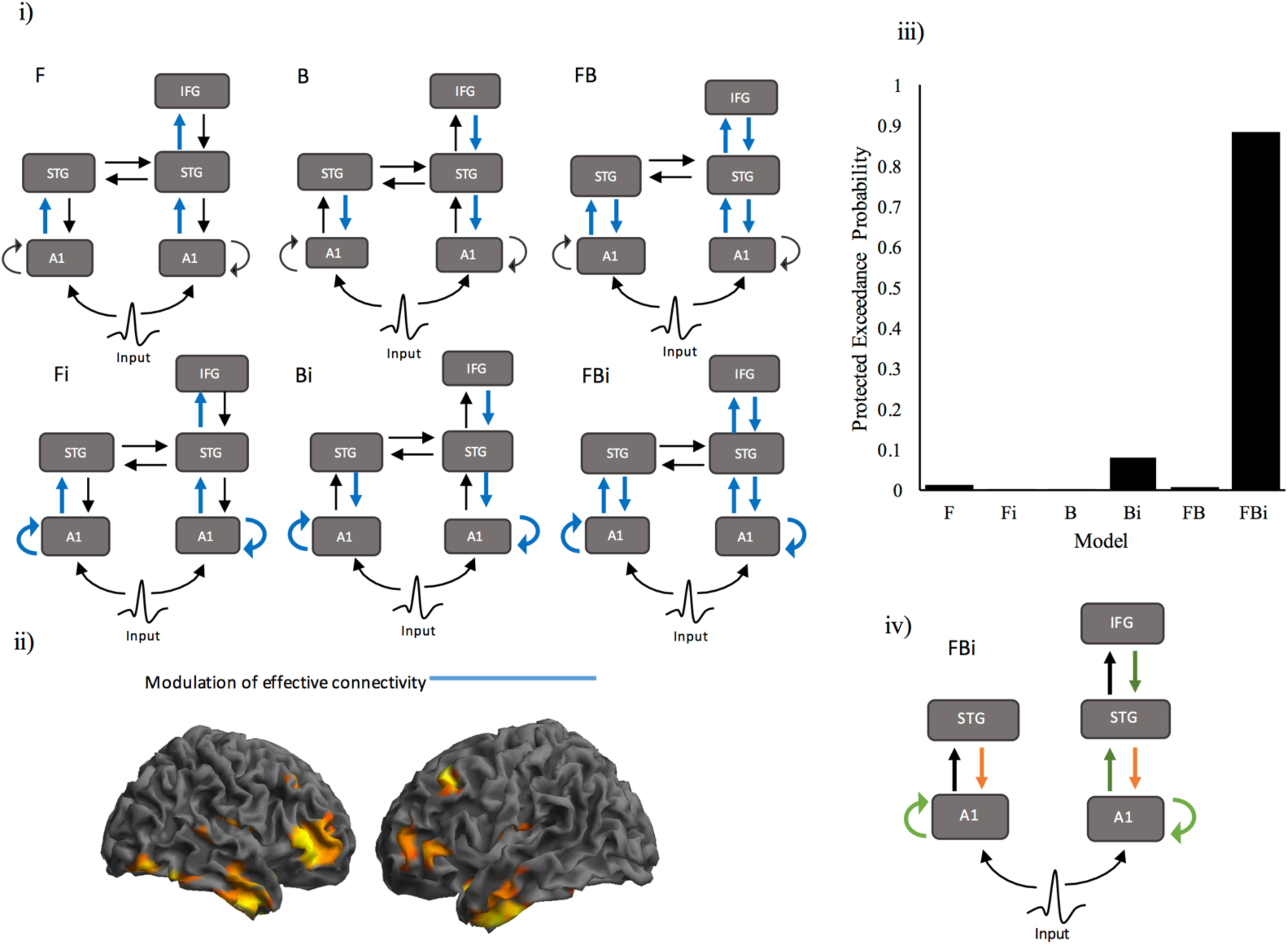
DCM specification and results for the MMN. i) The 6 models specified to assess the modulation of effective connectivity. This included three models of extrinsic modulation (forward (F), backward (B) and forward and backward (FB)) with intrinsic modulation in A1 either present (Fi, Bi, FBi) or absent. ii) Source localization statistical map for the *t*-test comparing deviant and standard tones, with clusters of significant voxels depicted in warm colours (*p*<.01 uncorrected). iii) Protected exceedance probabilities for the 6 models. BMS indicated that FBi was the winning model. iv) Posterior parameter estimates from the winning model. Green arrows depict connections significantly modulated by the presentation of the deviant. Orange arrows depict connections that differed in their modulation between the genetic groups.

#### 3.1.3. DCM of the Mismatch response

DCMs were specified to assess the modulation of extrinsic and intrinsic connectivity by the deviant tone. DCMs modelled a linear change in connectivity for tones 1 (deviant), 3, and 6 (standard). Tone 3 was included to ensure that the genetic differences apparent in the ERPs for repetition suppression were captured. Three different models of extrinsic modulation were examined: 1) forward, 2) backward, and 3) both forward and backward. Each of these were also combined with intrinsic modulation in A1 being either present or absent. This resulted in 6 models for comparison with BMS (Figure 4i). Subcortical input was modelled as entering the system through A1 bilaterally, with a post stimulus onset of 64ms. DCMs were modelled for a time-window from 0-400ms.

RFX BMS revealed that the model with the greatest model evidence was the full model, with forward and backward extrinsic modulation, as well as intrinsic modulation in A1 (Figure 4iii). We note that DCM corrects for the extra parameters in the full-model by accounting for the extra degrees of freedom introduced by extra free parameters. The PXP (0.98) provides strong evidence in favour of the winning model. The BOR (0.02) provide strong evidence that this result did not occur by chance. Classical inference on parameter estimates revealed a significant increase in left (*t*(39)=2.97, *p*=.005) and right (*t*(39)=3.02, p=.005) intrinsic A1 connectivity. Additionally, there was a marginal increase in connectivity from right A1 to right STG (*t*(39)=1.82, p=.076) as well as a marginal decrease in connectivity from right IFG to STG (*t*(39)=-1.85, *p*=.072) (Figure 4iv).

Finally, differences in network modulation between genotype groups were assessed using two sample *t*-tests on the parameter estimates (Figure 4iv). While there was an increase in STG to A1 backward connectivity for the deviant tone in Val homozygotes, Met carriers showed a decrease in these same connections. These differences were significant on the right (*t*(38)=2.19, *p*=.035) and marginal on the left (*t*(38)=1.80, *p*=.080).

### 3.2. LTP

#### 3.2.1. ERP analyses

The 2 x 2 ANOVA that probed the effect of time and tetanus showed an effect of time that confirmed potentiation had occurred. This included a left lateralized parieto-occipital time window from 132-156ms that peaked at 132ms (F_(2, 234)_ = 17.82, *p* < 0.001 FWE-c) consistent with the N170 (Figure 5i and i.a). In addition there was an occipital time window from 160-236ms that peaked at 188ms (F_(2, 234)_ = 28.86, *p* < 0.001 FWE-c) consistent with the P2 component (Figure 5ii and ii.a). There was also a small occipital peak at 92ms (F_(2, 234)_ = 28.86, *p* = 0.037 FWE-c) (Figure 5ii and ii.b). Additional significant clusters reflected the dipoles of both the N170 and P2. There was no significant effect of tetanus. There were also no significant interactions.

**Figure 5.**
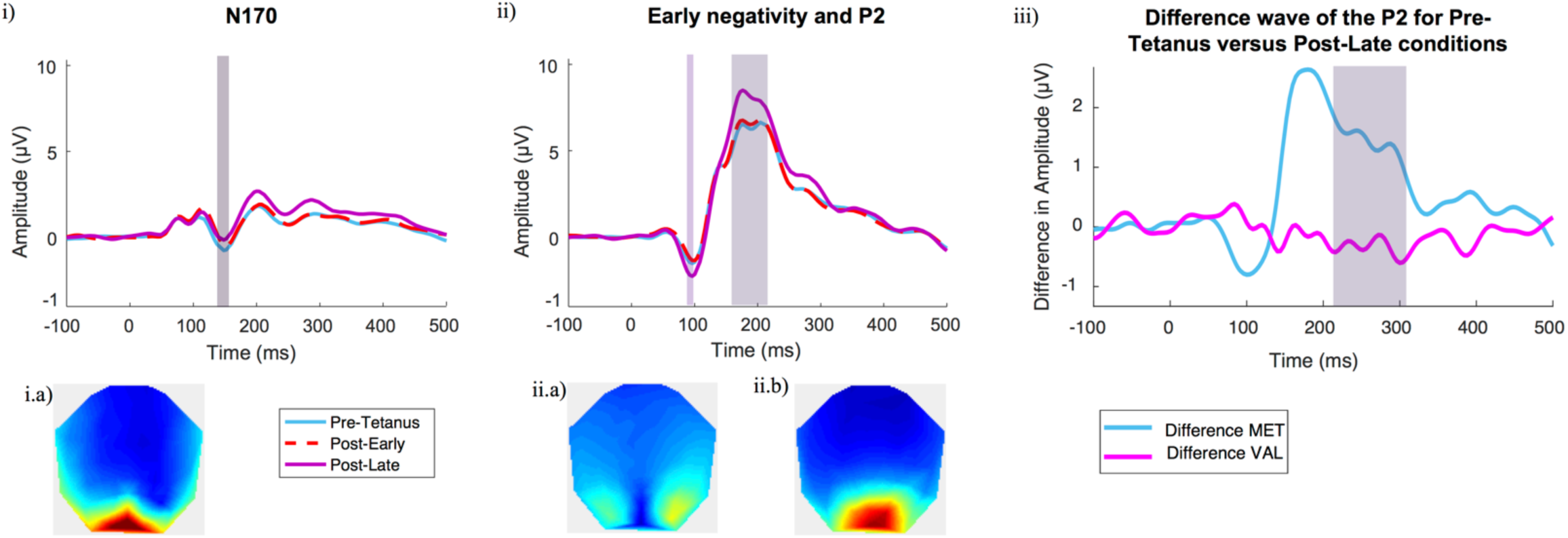
Averaged LTP ERPs across genotype i) at electrode P5, the ERP shows the small but significant decrease in the N170 component in the Post-Late condition. i.a) The topography of this component at 132ms. ii) Taken from electrode POz, there is a significant increase during Post-Late condition in the early negative deflection at 92ms and P2 component at 160-210ms. The respective topographies for each component are shown at i.a) 92ms and i.b) 188ms. iii) The difference in amplitude between Post-Late minus the Pre-Tetanus condition for Val homozygotes compared to Met carriers. As we found no effect of specificity for the tetanized stimulus, the graph shows the collapsed average of both tetanised and non-tetanised stimulus types.

Post-hoc contrasts for the effect of time revealed a change in an early (90ms) negative deflection and the N170 in both post tetanus conditions. A *t*-contrast confirmed this as an increase in negativity for the early negative deflection and decrease in negativity for the N170. In contrast, potentiation of the P2 was greatest in the post-late condition. A *t*-contrast confirmed this as an increase in positivity for the late condition compared to the early.

The 2x2x2 ANOVA that probed the effect of genotype, time post-tetanus and tetanizing stimulus showed that there was also a main effect of genotype that was approaching significance from 196-250ms and peaked at 248ms (F_(1, 152)_ = 18.41, *p* = 0.05) and 208ms (F_(1, 152)_ = 18.41, *p* = 0.07). Because of the highly conservative nature of the FWE-c this is interpreted as indicating a small effect at the P2. There was a main effect of time that replicated the effects of time reported above, comparing the early-post tetanus condition with the late post-tetanus condition. There was no main effect of tetanized stimulus. There were also no significant interactions. Post hoc contrasts of the effect of genotype showed that Met carriers had greater potentiation of the P2 component than Val homozygotes.

#### 3.2.2. Sources

Source analysis was performed on two time windows corresponding to the two significant clusters of activation from the ERP analysis (128-132ms and 188-208ms p<.05 uncorrected). There was substantial overlap in the sources for the two time windows, however the anterior sources in the 188-208ms time window were more robust. As such, this second time window was used to identify co-ordinates for Dynamic Causal Modelling (Figure 6ii). Previous literature has localized the potentiation of the N1b to V2/BA18 of the extrastriate visual cortex (Clapp et al., 2005; Teyler et al., 2005). Consistent with this, bilateral sources were identified in middle occipital gyrus (MOG, MNI coordinates left: -36 -90 4; right: 32, -92, 2). Additional significant sources were identified in left and right inferior temporal gyrus (ITG left: -52, -28, -24, right: 48, -12, -38) and left middle frontal gyrus (MFG, -26, 58, -4). This occipito-temporo-frontal network is consistent with networks important for visual memory (Miyashita, 1993) and memory consolidation (Laroche, Davis, & Jay, 2000).. These sources were then used for subsequent DCM analysis.

**Figure 6.**
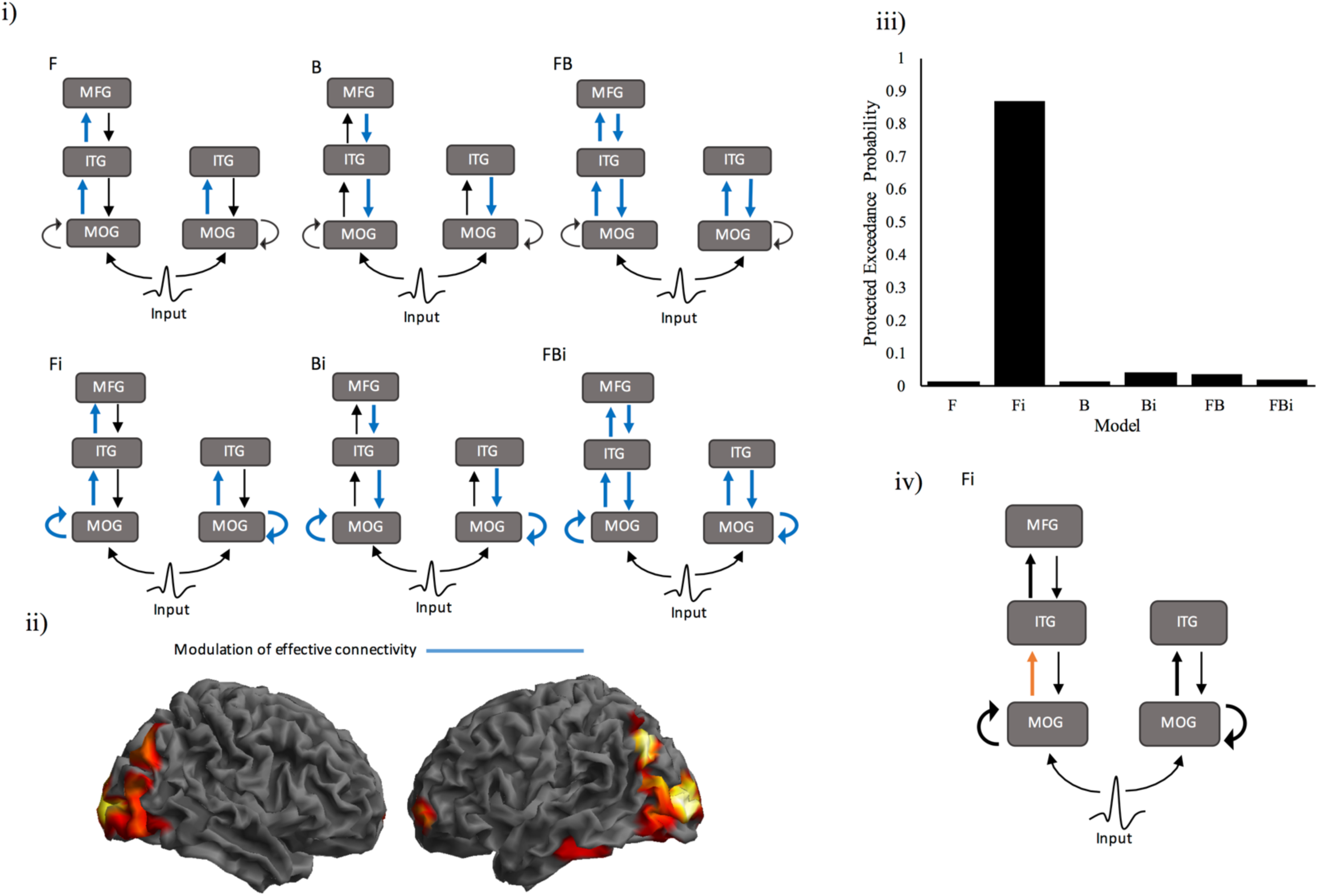
Specification and results for the DCMs modeling network modulation for the LTP paradigm. i) The 6 models specified to assess the modulation of effective connectivity. This included three models of extrinsic modulation (forward (F), backward (B) and forward and backward (FB)) with intrinsic modulation in MOG either present (Fi, Bi, FBi) or absent. ii) Source localization statistical map for the main effect of time, from the 3 (Time) x 2 (Tetanus) ANOVA for the 188-208ms time window, with clusters of significant voxels depicted in warm colours (*p*<.05 uncorrected). iii) Protected exceedance probabilities for the 6 models. BMS indicated that the Fi model was the winning model. iv) Posterior parameter estimates from the winning model. Green arrows depict connections significantly modulated by the paradigm.

#### 3.2.3. DCMs of Potentiation

Following an initial validation of connectivity between the sources of interest (see Supplementary Material), six DCMs were specified to assess the modulation of intrinsic and extrinsic connectivity following induction of LTP. Similar to the MMN paradigm, three models of extrinsic modulation were specified, in which the paradigm modulated 1) forward, 2) backward, or 3) forward and backward connectivity. These were each coupled with intrinsic modulation in MOG as either present or absent (Figure 6i). DCMs were specified for the time window from 0-350ms post stimulus presentation for the pre tetanus, early post tetanus, and late post tetanus blocks. Potentiation was modelled as a linear change in evoked response from pre-tetanus to early post-tetanus to late post-tetanus Input was modelled as entering the network through MOG with a post stimulus onset of 80ms.

BMS comparing the six models revealed that the model with the greatest model evidence was the model including modulation of forward extrinsic connections and intrinsic modulation within MOG (Figure 6ii). The PXP (0.87) provided strong evidence in favour of the winning model. The BOR (0.09) provided strong evidence that this result did not occur by chance.

*t*-tests on the parameter estimates revealed a significant increase in forward connectivity from left MOG to ITG (*t*(39) = 2.138, *p*=0.038) (Figure 6iii). However, there were no significant differences in network modulation between the two genotype groups.

## 4. Discussion

The aims of the current study were twofold. The first aim was to provide the first comparison between two electrophysiological paradigms designed to index sensory learning and plasticity. The Roving MMN paradigm has been widely used as an index of Predictive Coding mechanisms in the brain (Garrido, Kilner, Stephan, et al., 2009), while the visual LTP paradigm was designed to index Hebbian plasticity (Teyler et al., 2005). The second aim was to explore the differential sensitivity of the paradigms to group differences in evoked responses and underlying architecture. Groups were split based on participants’ expression of the *BDNF* Val66Met polymorphism. Brain Derived Neurotrophic Factor (BDNF) is an important mediator of synaptic plasticity (Tyler et al., 2002). As such, the *BDNF* polymorphism is implicated in the efficacy of NMDAR-dependent plasticity (Lamb et al., 2015), which plays a central role in both Predictive Coding and Hebbian plasticity.

Consistent with previous literature, the deviant tone in the roving MMN paradigm elicited a large fronto-central negativity peaking 172ms post stimulus (Garrido, Kilner, Stephan, et al., 2009; Näätänen, Paavilainen, Rinne, & Alho, 2007). Dynamic Causal Modelling (DCM) revealed that this response was generated by the modulation of primarily intrinsic, but also forward and backward extrinsic connectivity within a fronto-temporal network (Garrido et al., 2008, 2007). While there were no genetic differences in the amplitude of the MMN, *BDNF* Met carriers demonstrated reduced change between tones three and four in the tone sequences (indicating reduced variability), and reduced backward connectivity from STG to A1 compared to Val homozygotes. Also consistent with previous literature, ERP analysis of the visual LTP paradigm revealed a significant shift in the amplitude of the N1b in both the early and late post-tetanus conditions, as well as a shift in the P2a components of the VEP in the late post-tetanus block (Spriggs et al., 2017; Teyler et al., 2005). Within a generative network encompassing occipital, temporal and frontal sources, potentiation modulated forward extrinsic connectivity and intrinsic connectivity in middle occipital gyrus (MOG). While Met carriers showed greater potentiation of the P2 component of the VEP, there were no significant genetic differences in network modulation between the genetic groups. The current results therefore support the hypothesis that the paradigms index divergent processes underlying perceptual learning, which has important implications for future studies of aberrant plasticity in clinical populations.

### 4.1. MMN

The MMN ERP has a long history in electrophysiological research as an index of short term plasticity and cognitive function in healthy and clinical populations (Näätänen et al., 2007; Naatanen & Tiitinen, 2014). In recent years, the MMN has also been widely used as an index of perceptual inference under a Predictive Coding framework (Auksztulewicz & Friston, 2016; Garrido, Kilner, Stephan, et al., 2009). The current ERP results are consistent with the stereotypical expression of the MMN as a large fronto-central negativity, peaking 172ms after stimulus presentation. Additionally, the MMN response was found to be generated by modulations in forward, backward and intrinsic connectivity within the underlying fronto-temporal network. Classical inference on DCM parameter estimates revealed significant increases in intrinsic A1 connectivity, a marginal increase in forward connectivity from right A1 to right STG, and a marginal decrease in backward connectivity from right IFG to STG for the deviant tone. This is consistent with the predictive coding interpretation of MMN generation, under which the MMN is elicited by a disparity between sensory input and predictions that are made based on the memory trace from previous stimulation (Friston, 2005; Garrido, Kilner, Stephan, et al., 2009). A1 intrinsic connectivity is understood to represent the strength of memory formation, or more specifically, the precision of prediction errors. As this precision increases across successive presentations of a standard tone, it is inappropriately high for the presentation of a new deviant tone. Thus the deviant tone results in an increase in intrinsic A1 connectivity. This is coupled with an increase in bottom-up prediction error signals (due to the divergence between predictions and sensory input) and a decrease in the passing of inaccurate top-down predictions. These results are consistent with a large body of previous MMN studies that have identified similar network modulation for the deviant tone (Auksztulewicz & Friston, 2015; Boly et al., 2011; Garrido et al., 2008, 2007; Moran et al., 2014; Schmidt et al., 2013).

The current study also provides the first examination of the impact of the *BDNF* Val66Met polymorphism on repetition suppression and its underlying architecture using the roving MMN paradigm. Consistent with previous literature, there were no genotype differences in the amplitude of the MMN (Soltész et al., 2014), and thus no genetic difference in perceptual inference. However, Val homozygotes demonstrated greater variability than Met carriers in the difference waves of the ERP between tone presentations 3 and 4. More specifically, while Met carriers may have reached a ‘standard’ response by tone 3 and 4 (and thus there is less variability between the responses), Val homozygotes have not. This suggests that repetition suppression is faster in Met carriers, and thus indicates that perceptual learning is more efficient in this group.

Additionally, posterior parameter estimate extraction of the winning DCM revealed a decrease in backward connectivity from STG to A1 in *BDNF* Met carriers for the deviant tone. Conversely, Val homozygotes demonstrated an increase in these same connections. Suppression of predictions for the deviant tone has been widely documented in the IFG to STG connections of the MMN network (Boly et al., 2011; Garrido et al., 2008; Garrido, Kilner, Kiebel, et al., 2009). However, Met carriers demonstrate this suppression further down the processing hierarchy. As such, Met carriers may be better able to suppress inaccurate predictions at all levels of the processing hierarchy, which may lead to a faster repetition suppression (as revealed by the ERP analysis). Together, the current ERP and DCM results indicate that Met carriers may have an advantage when predictive coding mechanisms are employed.

### 4.2. LTP

The visual LTP paradigm was designed as a non-invasive parallel to the electrically-induced LTP protocols used with rodents. In this study, there was an enhancement of the P2 component of the VEP following high frequency, or ‘tetanic’, stimulation in the late-post tetanus block that is consistent with previous findings (Spriggs et al., 2017). There was also a significant enhancement of an early ERP component at around 90ms, which has also been found in one other study (Çavuş et al., 2012). These enhancements are understood to represent Hebbian plasticity processes within the visual cortex.

Interestingly, unlike in previous studies, there was no enhancement of the N1b component of the VEP following high frequency, or ‘tetanic’, stimulation (Clapp et al., 2005; McNair et al., 2006; Ross et al., 2008; Teyler et al., 2005). Instead we found a depressed response, or decrease in negativity, that was not only apparent immediately following the tetanus (early post-tetanus block), but remained present after a 30min break (late post-tetanus). It is not clear why this has occurred in our particular study, however, due to the established independence of the N170 and P2 peaks, it is not considered to confound interpretations of P2 potentiation (Crowley & Colrain, 2004; Freunberger, Klimesch, Doppelmayr, & Höller, 2007).

The visual system is highly hierarchical (Felleman & Van Essen, 1991; Salin & Bullier, 1995) and plasticity underlying visual perceptual learning begins at the earliest stages of visual processing (Furmanski, Schluppeck, & Engel, 2004; Masquelier & Thorpe, 2007; Schiltz et al., 1999; Schoups, Vogels, Qian, & Orban, 2001). The current results indicate that LTP induction not only modulates connectivity within early visual cortex, but also modulates forward connections between the striate/extrastriate, inferior temporal, and left prefrontal cortices. The inferior temporal lobe plays a crucial role in both object perception and visual memory (Miyashita, 1993). Occipito-temporal connections (corresponding to the ventral visual network) are thus central to perceptual learning, and rodent LTP has been extensively studied in both these regions (Artola & Singer, 1987; Berry, Teyler, & Taizhen, 1989; Bliss & Lømo, 1973; Heynen & Bear, 2001; Teyler & DiScenna, 1987). Additionally, LTP induction within the hippocampus has previously been shown to potentiate afferent connections to the prefrontal cortex (Gurden, Takita, & Jay, 2000; Jay, Burette, & Laroche, 1995; Laroche, Jay, & Thierry, 1990). These fronto-temporal connections are understood to be involved in memory consolidation and working memory (Laroche et al., 2000).

Classical inference on DCM posterior parameter estimates revealed a specific increase in forward connectivity from left visual cortex to inferior temporal gyrus (ITG) across the three time points (pre-tetanus, early post-tetanus and late post-tetanus). This increase indicates that there is an enhancement of the connectivity between these two regions following high frequency stimulation, and thus suggests that this pathway has undergone LTP. As opposed to the MMN, there was no modulation of backward connectivity. It is important to note that this does not suggest that backward connections are not present, rather it indicates that induction of LTP does not modulate these connections. Moreover, the absence of backward modulation suggests that the significance of the MFG source is not due to attentional modulation, and thus is more likely involved in memory consolidation.

The ERP analysis of the LTP paradigm revealed an effect of the *BDNF* Val66Met polymorphism on the P2 enhancement, where Met carriers demonstrated a greater increase in P2 amplitude than Val homozygotes. This is seemly contradictory to previous studies from our lab that have identified reduced potentiation of the N1b in Met carriers (Thompson et al., in prep). However, previous analyses have not looked at the P2 component. Additionally, the enhancement of the P2 component has not yet been comprehensively characterized, and it may represent a downstream consequence of the induction of LTP, or, as hypothesized by Spriggs et al., (2017), heterosynaptic LTD. However, there were no significant differences between the genotype groups in the modulation of forward or intrinsic connectivity of the underlying neuronal architecture.

### 4.3. Hebbian LTP & Predictive Coding

The current study presents the first comparison between two paradigms designed to index two different models of plasticity; Predictive Coding and Hebbian plasticity. Using the MMN as an index of Predictive Coding, and visual LTP as an index of Hebbian plasticity, Dynamic Causal Modelling revealed one principle difference between network modulations in the two paradigms: the generation of the MMN is dependent on the modulation of backward connectivity, while visually-induced LTP is not. Under Predictive Coding, backward connections are central to the generation of the evoked response, which represents the error between top-down (backwards) predictions and bottom-up sensory information (Garrido, Kilner, Stephan, et al., 2009; Huang & Rao, 2011). This study, and a number of previous studies, support this reciprocal relationship in the generation of the MMN (Auksztulewicz & Friston, 2015; Garrido et al., 2008, 2007; Moran et al., 2014; Schmidt et al., 2013). However, this does not appear to be the case for the LTP paradigm, where the winning DCM did not include modulation of backward connections. Under a Hebbian model, the strength of synaptic connections increases with repeated coactivation of neurons in a network (Cooke & Bliss, 2006; Teyler & DiScenna, 1987). The current results reflect this increase in forward connections only. As there is no need for comparison with a generative model, backward connections are no longer a driver in perceptual learning. The differences between Predictive Coding and LTP are depicted in Figure 7.

**Figure 7.**
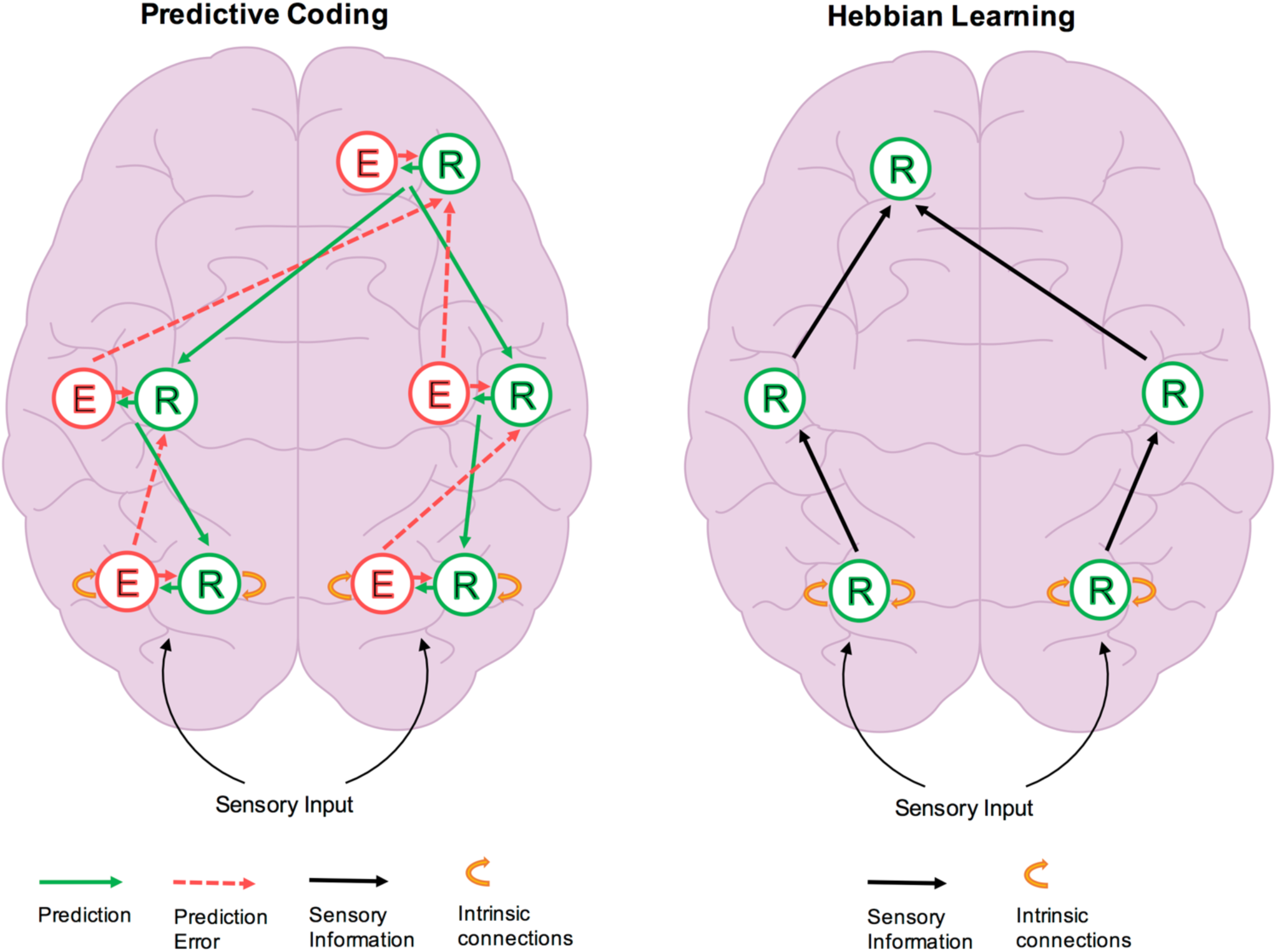
This figure shows a modified version of that presented by Stefanics, Kremláček, and Czigler (2014) to show the comparison between predictive coding and Hebbian learning forms of plasticity explored in this study. Under predictive coding, information passes between error units (E) and representation units (R). Backward connections carry predictions, whereas forward connections carry prediction errors (for example the MMN in response to a deviant tone). R units receive error information from the same node as well as lateral connections to nodes across the same level (not depicted) and lower hierarchical levels. Predictions (via backward connections are modulated and updated by the interaction between R and E units leading to repetition suppression or the standard response in the MMN task. Under Hebbian learning, representation units are updated and forward connections strengthened by the repetition of stimulus input, for example via photic tetanus in the visual LTP task.

The *BDNF* Val66Met polymorphism is understood to impact the efficacy of NMDAR-dependent plasticity (Chen et al., 2004; Egan et al., 2003; Lamb et al., 2015), which is central to both the generation of the MMN (Garrido, Kilner, Stephan, et al., 2009; Schmidt et al., 2013), and visual LTP (Clapp, Eckert, Teyler, & Abraham, 2006; Cooke & Bear, 2014). It would thus be reasonable to predict a comparable genetic group difference across both paradigms. However, genetic differences were only apparent in the modulation of backward connections for the MMN paradigm, with no significant genetic differences observed in the LTP generative network. This suggests that the *BDNF* polymorphism specifically modulates top-down message passing, which is central to Predictive Coding, but appears to be auxiliary for Hebbian mechanisms. It is important to note that the scalp level results indicate that there is a *BDNF*-mediated difference in the ERPs for the both paradigms, which suggests that there may be a *BDNF*-mediated effect that is not being fully captured by the LTP Dynamic Causal Models (discussed further below). Regardless, the current genetic results suggest that the sensory learning mechanisms indexed by the two paradigms differentially rely on NMDAR-mediated plasticity, thus supporting the hypothesis that they index different sensory encoding mechanisms.

The current results suggest that the mechanisms underlying experience-dependent sensory plasticity are not uniform across the brain and across different tasks. This is not to say that such mechanisms are unconditionally independent, and the Free Energy Principle demonstrates how Hebbian and Predictive Coding mechanisms can work in combination (Friston, 2005, 2009, 2010). Specifically, under the Free Energy Principle, biological agents aim to reduce surprise by minimizing free energy. Reducing free energy involves changing the ‘recognition density’ which is a probabilistic representation of the cause of sensory input (i.e., a generative model). Under the Predictive Coding framework, free energy is the difference between the recognition density and the sensory input, and is thus prediction error. Hebbian plasticity then optimizes the parameters of the generative model, encoding causal regularities. Therefore, the Free Energy Principle proposes complementary roles for Hebbian and Predictive Coding mechanisms in encoding hidden causes and states respectively. While not antagonistic to this hypothesis, the current results suggest that Hebbian mechanisms can also encode perceptual learning independent of a generative model. Further research will be required to determine how visual LTP is incorporated into a generative model.

Moreover, the current results do not suggest that the visual system is Hebbian and the auditory system is Bayesian. Numerous previous studies have identified examples of predictive coding within the visual system (e.g., Rao & Ballard, 1999; Rauss, Schwartz, & Pourtois, 2011), and the highly hierarchical and reciprocal nature of the visual system renders it the archetypal system for predictive mechanisms. What the current results do suggest is that different task demands elicit different encoding mechanisms. Exactly what ‘task demands’ elicit different encoding mechanisms is unclear, however, the primary difference between the paradigms is that the MMN is understood to result from short-term, echoic memory (Baldeweg, 2007), while LTP is the leading model of long-term memory (Cooke & Bliss, 2006). Again, this is consistent with the roles of Hebbian and Predictive Coding mechanisms encoding hidden causes and states respectively. It will be interesting for future studies to further characterize these differences.

Aberrant plasticity features in the neuropathology of a variety of psychological and neurological conditions from schizophrenia (Friston & Frith, 1995) to Alzheimer’s disease (Klein, 2006; Walsh, Drinkenburg, & Ahnaou, 2016). As such, the identification of disease related changes in plasticity has increasingly become a focus of electrophysiological research. The roving MMN and the visual-LTP paradigms have been independently used to demonstrate modulated plasticity in healthy and clinical populations (Boly et al., 2011; Dima et al., 2010; Normann et al., 2007; Rosch et al., 2017; Schmidt et al., 2013; Smallwood et al., 2015; Spriggs et al., 2017). However, the results of the current study indicate that the two paradigms not only measure different plasticity mechanisms, but also differ in the identification of group differences. This calls into question broad conclusions pertaining to the nature of ‘plasticity deficits’ underlying different disorders based on the results from one of these paradigms. Therefore, it may be beneficial for future studies exploring population differences in neuroplasticity to index multiple plasticity mechanisms, to ensure that any important differences between such mechanisms are not overlooked.

It is important to recognize a few limitations of the current study. Firstly, the reported posterior parameter estimates, and genetic effects on the ERPs are uncorrected for multiple comparisons. Primarily, this is due to the exploratory nature of these analyses. Secondly, the sample was largely female due to the involvement of the participants in an additional study. While it is not expected that there would be any sex differences in the measures collected, it will be important for future studies to recruit a more balanced cohort. Third, due to the low proportion of *BDNF* Met homozygotes in the general population (4-16%, Shimizu et al., 2004), Met homozygotes and heterozygotes were grouped into a single Met carrier group. While this is consistent with previous literature, it would be of interest to future studies to establish whether there is a dosage effect of the Met allele. Finally, as mentioned above, there were genetic differences in the ERPs for the LTP paradigm that did not manifest in the winning DCM. It is important to note that this is not due to an inadequate model fit (see Supplementary Material Figure S3). Instead, it may be that there are small but consistent genetic differences over multiple parameters, rather than a larger genetic effect on a small number of parameters. Thus it may be beneficial for future studies to take a more nuanced approach, potentially exploring different neuronal models, such as Canonical Microcircuits (Bastos et al., 2012; Moran, Pinotsis, & Friston, 2013), which may better capture any additional genetic differences in intrinsic connectivity.

The current study presents the first direct comparison between the visual LTP paradigm and auditory roving MMN paradigm in a cohort of healthy participants. While both paradigms are understood to index perceptual learning and plasticity, they are built on fundamentally different models of how experience dependent plasticity is encoded in the brain. In support of this, the current results indicate that the brain networks generating the LTP and MMN responses are modulated differently by the two paradigms. Additionally, the two paradigms are able to identify distinct group differences in the modulation of these brain networks. Therefore, the current study provides a demonstration of the heterogeneity of neural plasticity under differing task demands, and highlights the importance of comparison across paradigms when indexing modulated neuroplasticity in heathy and clinical populations.

## Acknowledgements

MJS is supported by Brain Research New Zealand Doctoral Scholarship. RLS is supported by Auckland Medical Research Foundation Doctoral scholarship. SDM is supported by the Rutherford Discovery Fellowship, Royal Society of New Zealand.

The authors would like to thank the Auckland Sequenom Facility for genetic analysis.

There are no conflicts of interest to report.

## References

Artola, A., & Singer, W. (1987). Long-term potentiation and NMDA receptors in rat visual cortex. Nature, 330(6149), 649–652.

Auksztulewicz, R., & Friston, K. (2015). Attentional enhancement of auditory mismatch responses: a DCM/MEG study. Cerebral Cortex, bhu323.

Auksztulewicz, R., & Friston, K. (2016). Repetition suppression and its contextual determinants in predictive coding. Cortex, 80, 125–140.

Baldeweg, T. (2007). ERP repetition effects and mismatch negativity generation: a predictive coding perspective. Journal of Psychophysiology, 21(3-4), 204–213.

Bastos, A. M., Usrey, W. M., Adams, R. A., Mangun, G. R., Fries, P., & Friston, K. J. (2012). Canonical microcircuits for predictive coding. Neuron, 76(4), 695–711.

Berry, R. L., Teyler, T. J., & Taizhen, H. (1989). Induction of LTP in rat primary visual cortex: tetanus parameters. Brain Research, 481(2), 221–227.

Bliss, T. V., & Lømo, T. (1973). Long-lasting potentiation of synaptic transmission in the dentate area of the anaesthetized rabbit following stimulation of the perforant path. The Journal of Physiology, 232(2), 331–356.

Boly, M., Garrido, M. I., Gosseries, O., Bruno, M.-A., Boveroux, P., Schnakers, C.,…Friston, K. (2011). Preserved feedforward but impaired top-down processes in the vegetative state. Science, 332(6031), 858–862.

Brainard, D. H. (1997). The psychophysics toolbox. Spatial Vision, 10, 433–436.

Çavuş, I., Reinhart, R. M., Roach, B. J., Gueorguieva, R., Teyler, T. J., Clapp, W. C.,…Mathalon, D. H. (2012). Impaired visual cortical plasticity in schizophrenia. Biological Psychiatry, 71(6), 512–520.

Chen, Z.-Y., Patel, P. D., Sant, G., Meng, C.-X., Teng, K. K., Hempstead, B. L., & Lee, F. S. (2004). Variant brain-derived neurotrophic factor (BDNF)(Met66) alters the intracellular trafficking and activity-dependent secretion of wild-type BDNF in neurosecretory cells and cortical neurons. Journal of Neuroscience, 24(18), 4401–4411.

Clapp, W. C., Eckert, M. J., Teyler, T. J., & Abraham, W. C. (2006). Rapid visual stimulation induces N-methyl-D-aspartate receptor-dependent sensory long-term potentiation in the rat cortex. Neuroreport, 17(5), 511–515.

Clapp, W. C., Hamm, J. P., Kirk, I. J., & Teyler, T. J. (2012). Translating long-term potentiation from animals to humans: a novel method for noninvasive assessment of cortical plasticity. Biological Psychiatry, 71(6), 496–502.

Clapp, W. C., Zaehle, T., Lutz, K., Marcar, V. L., Kirk, I. J., Hamm, J. P.,…Jancke, L. (2005). Effects of long-term potentiation in the human visual cortex: a functional magnetic resonance imaging study. Neuroreport, 16(18), 1977–1980.

Cooke, S. F., & Bear, M. F. (2014). How the mechanisms of long-term synaptic potentiation and depression serve experience-dependent plasticity in primary visual cortex. Phil. Trans. R. Soc. B, 369(1633), 20130284.

Cooke, S. F., & Bliss, T. V. P. (2006). Plasticity in the human central nervous system. Brain, 129(7), 1659–1673.

Corlett, P. R., Honey, G. D., & Fletcher, P. C. (2016). Prediction error, ketamine and psychosis: An updated model. Journal of Psychopharmacology, 269881116650087.

Crowley, K. E., & Colrain, I. M. (2004). A review of the evidence for P2 being an independent component process: age, sleep and modality. Clinical Neurophysiology, 115(4), 732–744.

D. O Hebb. (1949). The organization of behavior: a neuropsychological theory. New York: Wiley Science Editions.

David, O., Kiebel, S. J., Harrison, L. M., Mattout, J., Kilner, J. M., & Friston, K. J. (2006). Dynamic causal modeling of evoked responses in EEG and MEG. NeuroImage, 30(4), 1255–1272.

Desimone, R., Schein, S. J., Moran, J., & Ungerleider, L. G. (1985). Contour, color and shape analysis beyond the striate cortex. Vision Research, 25(3), 441–452.

Dima, D., Dietrich, D. E., Dillo, W., & Emrich, H. M. (2010). Impaired top-down processes in schizophrenia: A DCM study of ERPs. NeuroImage, 52(3), 824–832.

Egan, M. F., Kojima, M., Callicott, J. H., Goldberg, T. E., Kolachana, B. S., Bertolino, A.,…Weinberger, D. R. (2003). The BDNF val66met polymorphism affects activity-dependent secretion of BDNF and human memory and hippocampal function. Cell, 112(2), 257–269.

Fahle, M. (2004). Perceptual learning: A case for early selection. Journal of Vision, 4(10), 4–4.

Felleman, D. J., & Van Essen, D. C. (1991). Distributed hierarchical processing in the primate cerebral cortex. Cerebral Cortex, 1(1), 1–47.

Figurov, A., Pozzo-Miller, L. D., Olafsson, P., Wang, T., Lu B. (1996). Regulation of synaptic responses to high-frequency stimulation and LTP by neurotrophins in the hippocampus. Nature, 381(6584), 706.

Freunberger, R., Klimesch, W., Doppelmayr, M., & Höller, Y. (2007). Visual P2 component is related to theta phase-locking. Neuroscience Letters, 426(3), 181–186.

Friston, K. (2003). Learning and inference in the brain. Neural Networks, 16(9), 1325–1352.

Friston, K. (2005). A theory of cortical responses. Philosophical Transactions of the Royal Society of London B: Biological Sciences, 360(1456), 815–836.

Friston, K. (2009). The free-energy principle: a rough guide to the brain? Trends in Cognitive Sciences, 13(7), 293–301.

Friston, K. (2010). The free-energy principle: a unified brain theory? Nature Reviews Neuroscience, 11(2), 127–138.

Friston, K., Brown, H. R., Siemerkus, J., & Stephan, K. E. (2016). The dysconnection hypothesis (2016). Schizophrenia Research, 176(2), 83–94.

Friston, K. J. (2002). Bayesian estimation of dynamical systems: an application to fMRI. NeuroImage, 16(2), 513–530.

Friston, K. J., & Frith, C. D. (1995). Schizophrenia: a disconnection syndrome. Clin Neurosci, 3(2), 89–97.

Furmanski, C. S., Schluppeck, D., & Engel, S. A. (2004). Learning strengthens the response of primary visual cortex to simple patterns. Current Biology, 14(7), 573–578.

Garrido, M. I., Friston, K. J., Kiebel, S. J., Stephan, K. E., Baldeweg, T., & Kilner, J. M. (2008). The functional anatomy of the MMN: a DCM study of the roving paradigm. Neuroimage, 42(2), 936–944.

Garrido, M. I., Kilner, J. M., Kiebel, S. J., Stephan, K. E., Baldeweg, T., & Friston, K. J. (2009). Repetition suppression and plasticity in the human brain. Neuroimage, 48(1), 269–279.

Garrido, M. I., Kilner, J. M., Kiebel, S. J., Stephan, K. E., & Friston, K. J. (2007). Dynamic causal modelling of evoked potentials: a reproducibility study. Neuroimage, 36(3), 571–580.

Garrido, M. I., Kilner, J. M., Stephan, K. E., & Friston, K. J. (2009). The mismatch negativity: a review of underlying mechanisms. Clinical Neurophysiology, 120(3), 453–463.

Goldberg, T. E., & Weinberger, D. R. (2004). Genes and the parsing of cognitive processes. Trends in Cognitive Sciences, 8(7), 325–335.

Grill-Spector, K., & Malach, R. (2004). The human visual cortex. Annual Reviews Neuroscience, 27, 649–677.

Gurden, H., Takita, M., & Jay, T. M. (2000). Essential role of D1 but not D2 receptors in the NMDA receptor-dependent long-term potentiation at hippocampal-prefrontal cortex synapses in vivo. The Journal of Neuroscience: The Official Journal of the Society for Neuroscience, 20(22), RC106–RC106.

Hariri, A. R., Goldberg, T. E., Mattay, V. S., Kolachana, B. S., Callicott, J. H., Egan, M. F., & Weinberger, D. R. (2003). Brain-derived neurotrophic factor val66met polymorphism affects human memory-related hippocampal activity and predicts memory performance. Journal of Neuroscience, 23(17), 6690–6694.

Harris, E. W., Ganong, A. H., & Cotman, C. W. (1984). Long-term potentiation in the hippocampus involves activation of N-methyl-D-aspartate receptors. Brain Research, 323(1), 132–137.

Heynen, A. J., & Bear, M. F. (2001). Long-term potentiation of thalamocortical transmission in the adult visual cortex in vivo. Journal of Neuroscience, 21(24), 9801–9813.

Huang, Y., & Rao, R. P. (2011). Predictive coding. Wiley Interdisciplinary Reviews: Cognitive Science, 2(5), 580–593.

Jansen, B. H., & Rit, V. G. (1995). Electroencephalogram and visual evoked potential generation in a mathematical model of coupled cortical columns. Biological Cybernetics, 73(4), 357–366.

Jay, T. M., Burette, F., & Laroche, S. (1995). NMDA Receptor-dependent Long-term Potentiation in the Hippocampal Afferent Fibre System to the Prefrontal Cortex in the Rat. European Journal of Neuroscience, 7(2), 247–250.

Kambeitz, J. P., Bhattacharyya, S., Kambeitz-Ilankovic, L. M., Valli, I., Collier, D. A., & McGuire, P. (2012). Effect of BDNF val 66 met polymorphism on declarative memory and its neural substrate: a meta-analysis. Neuroscience & Biobehavioral Reviews, 36(9), 2165–2177.

Kiebel, S. J., David, O., & Friston, K. J. (2006). Dynamic causal modelling of evoked responses in EEG/MEG with lead field parameterization. NeuroImage, 30(4), 1273–1284.

Kirk, I. J., McNair, N. A., Hamm, J. P., Clapp, W. C., Mathalon, D. H., Cavus, I., & Teyler, T. J. (2010). Long-term potentiation (LTP) of human sensory-evoked potentials. Wiley Interdisciplinary Reviews: Cognitive Science, 1(5), 766–773.

Kirkwood, A., & Bear, M. F. (1994). Hebbian synapses in visual cortex. Journal of Neuroscience, 14(3), 1634–1645.

Klein, W. L. (2006). Synaptic targeting by Aβ oligomers (ADDLS) as a basis for memory loss in early Alzheimer’s disease. Alzheimer’s & Dementia, 2(1), 43–55.

Kleiner, M., Brainard, D., Pelli, D., Ingling, A., Murray, R., Broussard, C., & others. (2007). What’s new in Psychtoolbox-3. Perception, 36(14), 1.

Knill, D. C., & Pouget, A. (2004). The Bayesian brain: the role of uncertainty in neural coding and computation. Trends in Neurosciences, 27(12), 712–719.

Koch, C., & Poggio, T. (1999). Predicting the visual world: silence is golden. Nature Neuroscience, 2, 9–10.

Kourtzi, Z., & DiCarlo, J. J. (2006). Learning and neural plasticity in visual object recognition. Current Opinion in Neurobiology, 16(2), 152–158.

Lamb, Y. N., McKay, N. S., Thompson, C. S., Hamm, J. P., Waldie, K. E., & Kirk, I. J. (2015). Brain-derived neurotrophic factor Val66Met polymorphism, human memory, and synaptic neuroplasticity. Wiley Interdisciplinary Reviews: Cognitive Science, 6(2), 97–108.

Laroche, S., Davis, S., & Jay, T. M. (2000). Plasticity at hippocampal to prefrontal cortex synapses: dual roles in working memory and consolidation. Hippocampus, 10(4), 438–446.

Laroche, S., Jay, T. M., & Thierry, A.-M. (1990). Long-term potentiation in the prefrontal cortex following stimulation of the hippocampal CA1/subicular region. Neuroscience Letters, 114(2), 184–190.

Litvak, V., & Friston, K. (2008). Electromagnetic source reconstruction for group studies. Neuroimage, 42(4), 1490–1498.

Masquelier, T., & Thorpe, S. J. (2007). Unsupervised learning of visual features through spike timing dependent plasticity. PLoS Computational Biology, 3(2), e31.

McNair, N. A., Clapp, W. C., Hamm, J. P., Teyler, T. J., Corballis, M. C., & Kirk, I. J. (2006). Spatial frequency-specific potentiation of human visual-evoked potentials. Neuroreport, 17(7), 739–741.

Miyashita, Y. (1993). Inferior temporal cortex: where visual perception meets memory. Annual Review of Neuroscience, 16(1), 245–263.

Moran, R. J., Campo, P., Symmonds, M., Stephan, K. E., Dolan, R. J., & Friston, K. J. (2013). Free energy, precision and learning: the role of cholinergic neuromodulation. Journal of Neuroscience, 33(19), 8227–8236.

Moran, R. J., Symmonds, M., Dolan, R. J., & Friston, K. J. (2014). The brain ages optimally to model its environment: evidence from sensory learning over the adult lifespan. PLoS Computational Biology, 10(1), e1003422.

Moran, R., Pinotsis, D. A., & Friston, K. (2013). Neural masses and fields in dynamic causal modeling. Frontiers in Computational Neuroscience, 7. Retrieved from https://www.ncbi.nlm.nih.gov/pmc/articles/PMC3664834/

Muthukumaraswamy, S. D., Shaw, A. D., Jackson, L. E., Hall, J., Moran, R., & Saxena, N. (2015). Evidence that subanesthetic doses of ketamine cause sustained disruptions of NMDA and AMPA-mediated frontoparietal connectivity in humans. Journal of Neuroscience, 35(33), 11694–11706.

Näätänen, R., Paavilainen, P., Rinne, T., & Alho, K. (2007). The mismatch negativity (MMN) in basic research of central auditory processing: a review. Clinical Neurophysiology, 118(12), 2544–2590.

Naatanen, R., & Tiitinen, H. (2014). Auditory information processing as indexed by the mismatch negativity. Advances in Psychological Science Biological and Cognitive Aspects, 145–170.

Normann, C., Schmitz, D., Fürmaier, A., Döing, C., & Bach, M. (2007). Long-term plasticity of visually evoked potentials in humans is altered in major depression. Biological Psychiatry, 62(5), 373–380.

Park, H., & Poo, M. (2013). Neurotrophin regulation of neural circuit development and function. Nature Reviews Neuroscience, 14(1), 7–23.

Pelli, D. G. (1997). The VideoToolbox software for visual psychophysics: Transforming numbers into movies. Spatial Vision, 10(4), 437–442.

Rao, R. P., & Ballard, D. H. (1999). Predictive coding in the visual cortex: a functional interpretation of some extra-classical receptive-field effects. Nature Neuroscience, 2(1), 79–87.

Rauss, K., Schwartz, S., & Pourtois, G. (2011). Top-down effects on early visual processing in humans: A predictive coding framework. Neuroscience & Biobehavioral Reviews, 35(5), 1237–1253.

Rigoux, L., Stephan, K. E., Friston, K. J., & Daunizeau, J. (2014). Bayesian model selection for group studies–revisited. Neuroimage, 84, 971–985.

Rinne, T., Alho, K., Ilmoniemi, R. J., Virtanen, J., & Näätänen, R. (2000). Separate time behaviors of the temporal and frontal mismatch negativity sources. Neuroimage, 12(1), 14–19.

Rosch, R. E., Auksztulewicz, R., Leung, P. D., Friston, K. J., & Baldeweg, T. (2017). NMDA receptor blockade causes selective prefrontal disinhibition in a roving auditory oddball paradigm. bioRxiv, 133371.

Ross, R. M., McNair, N. A., Fairhall, S. L., Clapp, W. C., Hamm, J. P., Teyler, T. J., & Kirk, I. J. (2008). Induction of orientation-specific LTP-like changes in human visual evoked potentials by rapid sensory stimulation. Brain Research Bulletin, 76(1), 97–101.

Salin, P.-A., & Bullier, J. (1995). Corticocortical connections in the visual system: structure and function. Physiological Reviews, 75(1), 107–155.

Schiltz, C., Bodart, J. M., Dubois, S., Dejardin, S., Michel, C., Roucoux, A.,…Orban, G. A. (1999). Neuronal mechanisms of perceptual learning: changes in human brain activity with training in orientation discrimination. Neuroimage, 9(1), 46–62.

Schmidt, A., Diaconescu, A. O., Kometer, M., Friston, K. J., Stephan, K. E., & Vollenweider, F. X. (2013). Modeling ketamine effects on synaptic plasticity during the mismatch negativity. Cerebral Cortex, 23(10), 2394–2406.

Schoups, A., Vogels, R., Qian, N., & Orban, G. (2001). Practising orientation identification improves orientation coding in V1 neurons. Nature, 412(6846), 549–553.

Shimizu, E., Hashimoto, K., & Iyo, M. (2004). Ethnic difference of the BDNF 196G/A (val66met) polymorphism frequencies: the possibility to explain ethnic mental traits. American Journal of Medical Genetics Part B: Neuropsychiatric Genetics, 126(1), 122–123.

Smallwood, N., Spriggs, M. J., Thompson, C. S., Wu, C. C., Hamm, J. P., Moreau, D., & Kirk, I. J. (2015). Influence of physical activity on human sensory long-term potentiation. Journal of the International Neuropsychological Society, 21(10), 831–840.

Soltész, F., Suckling, J., Lawrence, P., Tait, R., Ooi, C., Bentley, G.,…others. (2014). Identification of BDNF sensitive electrophysiological markers of synaptic activity and their structural correlates in healthy subjects using a genetic approach utilizing the functional BDNF Val66Met polymorphism. PloS One, 9(4), e95558.

Spriggs, M. J., Cadwallader, C. J., Hamm, J. P., Tippett, L. J., & Kirk, I. J. (2017). Age-related alterations in human neocortical plasticity. Brain Research Bulletin, 130, 53–59.

Stefanics, G., Kremláček, J., & Czigler, I. (2014). Visual mismatch negativity: a predictive coding view. Frontiers in Human Neuroscience, 8. Retrieved from https://www.ncbi.nlm.nih.gov/pmc/articles/PMC4165279/

Teyler, T. J., & DiScenna, P. (1987). Long-term potentiation. Annual Review of Neuroscience, 10(1), 131–161.

Teyler, T. J., Hamm, J. P., Clapp, W. C., Johnson, B. W., Corballis, M. C., & Kirk, I. J. (2005). Long-term potentiation of human visual evoked responses. European Journal of Neuroscience, 21(7), 2045–2050.

Tyler, W. J., Alonso, M., Bramham, C. R., & Pozzo-Miller, L. D. (2002). From acquisition to consolidation: on the role of brain-derived neurotrophic factor signaling in hippocampal-dependent learning. Learning & Memory, 9(5), 224–237.

Walsh, C., Drinkenburg, W., & Ahnaou, A. (2016). Neurophysiological Assessment of Neural Network Plasticity and Connectivity: Progress towards Early Functional Biomarkers for Disease Interception Therapies in Alzheimer’s disease. Neuroscience & Biobehavioral Reviews. Retrieved from http://www.sciencedirect.com.ezproxy.auckland.ac.nz/science/article/pii/S014976341_6304456

